# Resolving Cytosolic Diffusive States in Bacteria by Single-Molecule Tracking

**DOI:** 10.1101/483321

**Authors:** J. Rocha, J. Corbitt, T. Yan, C. Richardson, A. Gahlmann

## Abstract

The trajectory of a single protein in the cytosol of a living cell contains information about its molecular interactions in its native environment. However, it has remained challenging to accurately resolve and characterize the diffusive states that can manifest in the cytosol using analytical approaches based on simplifying assumptions. Here, we show that multiple intracellular diffusive states can be successfully resolved if sufficient single-molecule trajectory information is available to generate well-sampled distributions of experimental measurements and if experimental biases are taken into account during data analysis. To address the inherent experimental biases in camera-based and MINFLUX-based single-molecule tracking, we use an empirical data analysis framework based on Monte Carlo simulations of confined Brownian motion. This framework is general and adaptable to arbitrary cell geometries and data acquisition parameters employed in 2D or 3D single-molecule tracking. We show that, in addition to determining the diffusion coefficients and populations of prevalent diffusive states, the timescales of diffusive state switching can be determined by stepwise increasing the time window of averaging over subsequent single-molecule displacements. Time-averaged diffusion (TAD) analysis of single-molecule tracking data may thus provide quantitative insights into binding and unbinding reactions among rapidly diffusing molecules that are integral for cellular functions.

## Introduction

The ability to probe the positions and motions of single molecules in living cells has made single-molecule localization and tracking microscopy a powerful experimental tool to study the molecular basis of cellular functions (1-3). Single-molecule trajectories, if sampled in sufficient numbers, provide the distribution of molecular motion behavior in cells, and statistical analyses of localization and trajectory data has been used to resolve the prevalent diffusive states as well as their population fractions. A key benefit of tracking single molecules is that individual trajectories can be sorted according to predefined (quality) metrics, for example, to include only non-blinking molecules (4), or molecules localized in specific subcellular regions of interest (5). These advantages are not shared by ensemble-averaged measurements such as fluorescence recovery after photobleaching (FRAP) and fluorescence correlation spectroscopy (FCS) (6).

Bacteria are ideally suited specimens for single-molecule localization and tracking microscopy. Unlike eukaryotic cells, the small size of bacteria (~1 μm in diameter) guarantees that all molecules remain in focus during imaging (7), particularly when the microscope uses an engineered 3D point-spread-function (PSF), such as an astigmatic (8) or a double-helix PSF (DHPSF) (9, 10). Early applications of single-molecule localization microscopy in bacteria focused on differentiating stationary vs. freely diffusing molecules and quantifying the relative population fractions and lifetimes of these diffusive states. For example, DNA bound lac repressors in search of their promoter region appear stationary at 10 ms frame rates and can thus be clearly distinguished from unbound lac repressors which explore the entire *E. coli* cell volume on the same timescale (11). Similarly, the *E. coli* chromosome-partitioning protein MukB forms stationary clusters only when incorporated into the quasi-static DNA-bound structural maintenance of chromosomes (SMC) complex (12). In both of these cases, the stationary, DNA-bound states represent the biologically active form of the protein while the unbound diffusive state represents the inactive protein. However, other proteins, in particular those involved in delocalized regulatory and signaling networks, may not exhibit such stationary states. These proteins may instead form oligomeric complexes that diffuse at measurably different rates (13-17). A major objective for single-molecule tracking microscopy is therefore to resolve the different diffusive states that manifest in the cytosol of living cells.

Assigning a single molecule to a specific diffusive state is challenging, especially for fast diffusing cytosolic species. The molecular displacements measured in single-molecule tracking can be used to compute apparent diffusion coefficients for each detected single molecule, but these estimates are prone to large errors, particularly when the trajectories are short and the number of available molecular displacements are low (15, 18). Short trajectories (<20 displacements) are the norm in live-cell single-molecule tracking with genetically encodable fluorescent protein labels. However, genetically encoded fluorescent proteins offer unmatched labeling specificity and efficiency and therefore remain preferable when off-target labeling with chemical dyes may lead to artifacts (19). For slowly diffusing molecules in bacteria, it is possible to resolve multiple diffusive states by fitting the experimentally measured distributions of molecular displacements, *r*, or apparent diffusion coefficients, *D\**, using analytical equations describing Brownian, i.e. normal, diffusion (15, 18, 20-22). Such analytical approaches produce acceptable results only if biomolecular motion is slow enough that confinement effects can be ignored. However, a typical cytosolic protein undergoing Brownian diffusion at a rate *D* = 10 μm^2^/s can traverse the entire width of a rod-shaped bacterial cell in as little as 10-25 milliseconds. As a result, observed motion of cytosolic proteins in bacteria is strongly confined by the cell boundaries and molecular displacements will, on average, be smaller than those expected for unconfined diffusion. Approaches assuming unconfined Brownian motion are therefore not suitable when tracking fast diffusing molecules in the cytosol of bacterial cells.

Several approaches have been developed in recent years to extract the diffusion rates and population fractions of different diffusive states that manifest for unbound molecules in confined cellular environments. These approaches account for confinement effects by the cell boundaries either (semi-)analytically (23-26) or numerically through Monte Carlo simulation of Brownian diffusion trajectories (7, 13, 17, 27, 28). Here, we test and experimentally validate a numerical analysis framework based on Monte Carlo simulations for both 2D and 3D single-molecule tracking in bacterial cells (**Fig. 1**). By explicitly accounting for confinement as well as ‘motion-blur’ of diffusing molecules inside small bacterial cells, we extract the *unconfined* diffusion coefficients for two genetically encoded fluorescence proteins, eYFP and mEos3.2, in living *Y. enterocolitica* cells. Using simulated 2D or 3D single-molecule tracking data of known diffusive state composition, we quantify to what extent two or more simultaneously present diffusive states can be resolved by numerical fitting of the displacement or apparent diffusion coefficient distributions. Finally, we consider the influence of dynamic transitions between different diffusive states that may manifest upon association and dissociation of freely diffusing molecules. We propose a new approach, based on time-averaged diffusion (TAD) analysis, to determine the timescales of such association and dissociation dynamics. We conclude that quantitative numerical analysis of 2D and 3D single-molecule trajectories can provide accurate estimations of diffusion rates, population fractions, and interconversion rates of prevalent intracellular diffusive states. Such information is crucial for investigating the dynamic molecular-level events that regulate the functional outputs of signaling and control networks in living cells.

**Figure 1.**
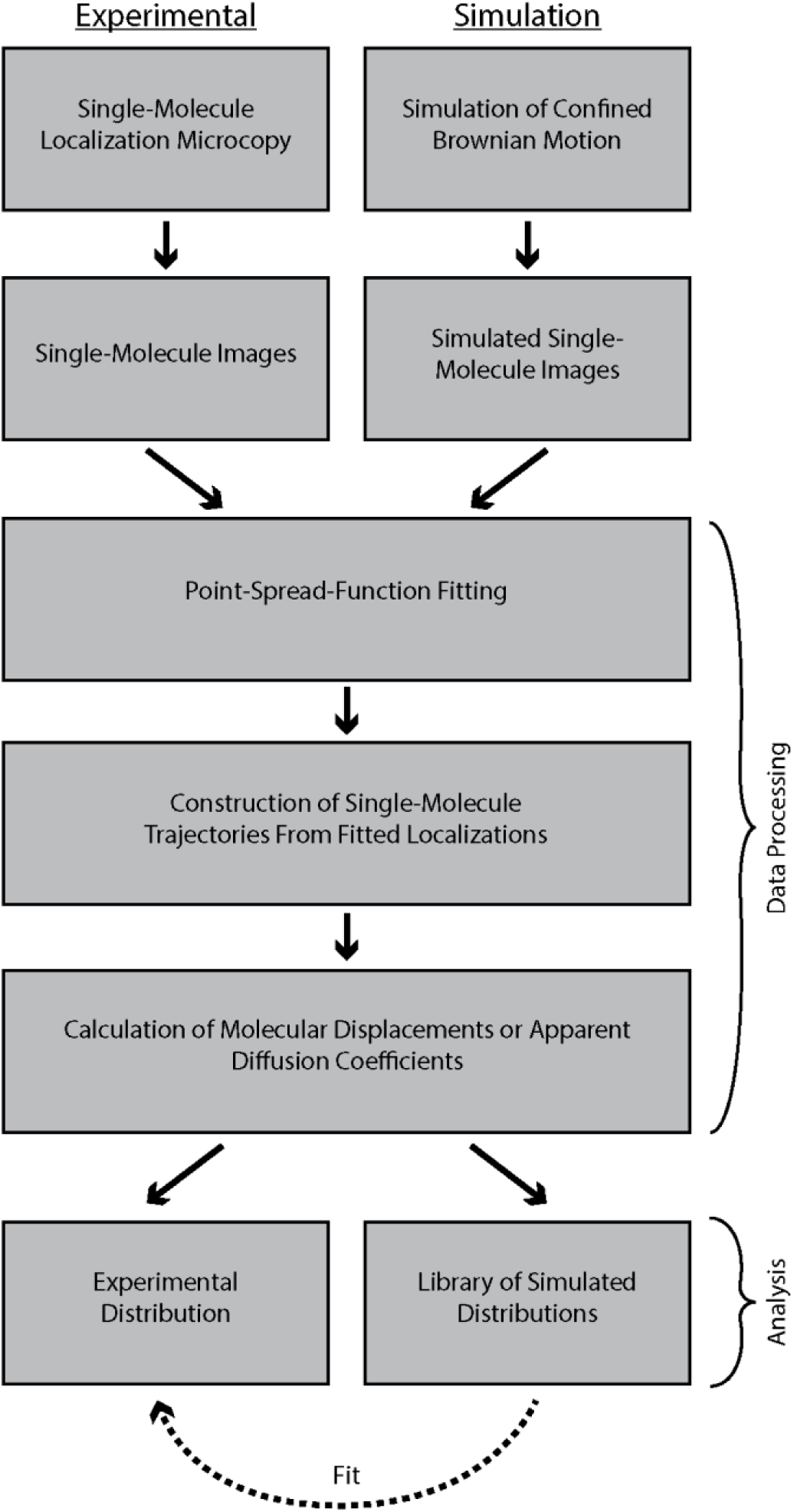
Diagram of numerical diffusion fitting analysis workflow. Experimental and simulated data are analyzed using the same data processing routines so that experimentally determined apparent diffusion coefficient (or displacement) distributions can be analyzed using linear combinations of simulated distributions.

## Materials and Methods

### Super-resolution Fluorescence Imaging Setup

Experiments were performed on a custom-built dual-color inverted fluorescence microscope based on the RM21 platform (Mad City Labs, Inc, Madison, Wisconsin). Immersion oil was placed between the objective lens (UPLSAPO 100X 1.4 NA) and the glass cover slip (VWR, Radnor, Pennsylvania, #1.5, 22mmx22mm). A 514 nm laser (Coherent, Santa Clara, California, Genesis MX514 MTM) was used for excitation of eYFP (~350 W/cm^2^) and 561 nm laser (Coherent Genesis MX561 MTM) was used for excitation of mEos3.2 (~350 W/cm^2^). A 405 nm laser (Coherent OBIS 405nm LX) was used to activate mEos3.2 (~20 W/cm^2^) simultaneously with 561nm excitation. Single-molecule images were obtained by utilizing eYFP photoblinking (29) and mEos3.2 photo-switching. Zero-order quarter-wave plates (Thorlabs, Newton, New Jersey, WPQ05M-405, WPQ05M-514, WPQ05M-561) were used to circularly polarize all excitation lasers. The spectral profile of the 514nm laser was filtered using a bandpass filter (Chroma, Bellows Falls, Vermont, ET510/10bp). Fluorescence emission was passed through a shared filter set (Semrock, Rochester, New York, LP02-514RU-25, Semrock NF03-561E-25, and Chroma ET700SP-2P8). A dichroic beam splitter (Chroma T560lpxr-uf3) was then used to split the emission pathway into ‘green’ and ‘red’ channels to image eYFP and mEos3.2, respectively. An additional 561nm notch filter (Chroma ZET561NF) was inserted into the ‘red’ channel to block scattered laser light. Each emission path contains a wavelength-specific dielectric phase mask (Double Helix, LLC, Boulder, Colorado) that is placed in the Fourier plane of the microscope to generate a DHPSF (10, 30). The fluorescence signals in both channels are detected on two separate sCMOS cameras (Hamamatsu, Bridgewater, New Jersey, ORCA-Flash 4.0 V2). Up to 20,000 frames are collected per field-of-view with an exposure time of 25ms. Exposure times of 25ms were used for all experiments to maximize fluorescent signal to background ratio (31). A flip-mirror in the emission pathway enables toggling the microscope between fluorescence imaging and phase contrast imaging modes without having to change the objective lens of the microscope.

### Raw Data Processing

Raw single-molecule PSF images were processed and analyzed using MATLAB (The MathWorks, Inc, Natick, Massachusetts). Standard PSF images were analyzed using centroid estimation (32). DHPSF images were analyzed using a modified version of the easyDHPSF code (33). Specifically, maximum likelihood estimation based on a double-Gaussian PSF model was used to extract the 3D localizations of single-molecule emitters (34). For experimental data, the background was estimated using a median filter with a time window of 10 frames (35).

To assign localizations to individual cells, cell outlines were generated based on the phase contrast images using the open-source software OUFTI (36). The cell outlines were transformed to overlay on the fluorescence data by a two-step 2D affine transformation using the ‘cp2tform’ function in MATLAB. First, five control point pairs were manually selected by estimating the position of the cell poles of the same five cells in both the single-molecule localization data and cell outlines. A rough transformation was generated, and cell outlines containing less than 10 localizations within their boundaries were removed. In addition, cells positioned partly outside the field-of-view were manually removed so they do not skew the final transformation. The center of mass for all remaining cell outlines and single-molecule localizations within them then served as a larger set of control point pairs to compute the final transformation function. Only localizations that lie within the cell outlines after transformation were considered for further analysis.

### Single Molecule Tracking Analysis

Molecular displacements were computed as the Euclidean distance between subsequent localizations of the same molecule using a distance threshold of 2.5 μm. Displacements were linked into a trajectories and considered for further analysis only if at least 3 subsequent (i.e. localizations in adjacent frames) displacements were available. In addition, if two or more localizations were present in the cell simultaneously during the length of the trajectory, the trajectory was discarded. These steps minimized miss-assignment of two or more molecules to the same trajectory (37).

To obtain apparent diffusion coefficients for a given trajectory, its Mean Squared Displacement (MSD) was calculated using

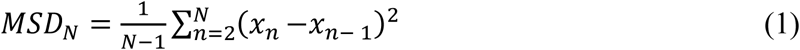

where *N* is the total number of localizations in the trajectory and *x*_*n*_ is the position of the molecule at time point *n*. The apparent diffusion coefficient, *D\** was then computed by

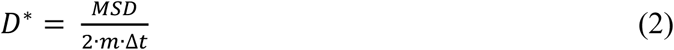

where *m* = 2 or 3 is the dimensionality and *Δt* =25 ms is the camera exposure time used in all our experiments and simulations. We note that the so-estimated single-step apparent diffusion coefficients and displacements do not directly take into account static and dynamic localization errors (18), or the effect of confinement within the bacterial cells. We instead account for these effects through explicit simulation of experimental data, as described in the following section.

### Monte Carlo Simulations for Camera-Based Tracking

Calculation of the apparent diffusion coefficients for a large number of tracked molecules will result in a distribution of values even if molecular diffusion is governed by a single diffusive state. In addition, for confined diffusion within small bacterial cell volumes, the movement of molecules is restricted in space. Such confinement results in an overall left shift of the apparent diffusion coefficient distributions for a given diffusive state (**Fig 2a**, dashed lines). The shape of the confined distribution is dependent on the size and shape of the confining volume.

**Figure 2.**
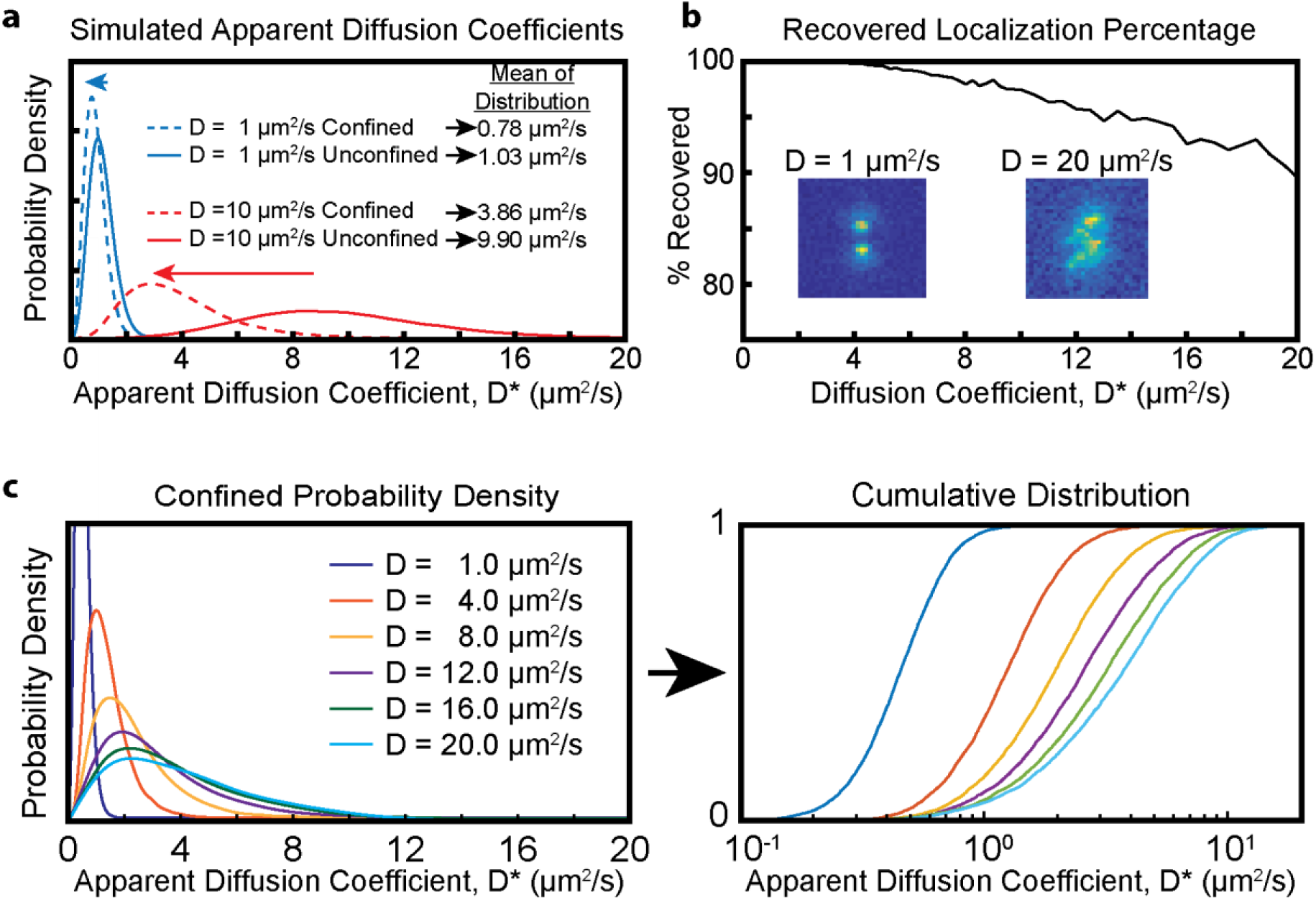
Monte-Carlo simulations of expected experimental distribution. (a) Probability density functions showing the effect of spatial confinement. The apparent diffusion coefficients are computed based on the time-integrated (25 ms) center-of-mass coordinates of simulated particles undergoing Brownian diffusion in a cylindrical volume (radius = 0.4 μm, length = 5 μm). The confined distributions are left-shifted (dashed lines) compared to the unconfined distributions. (b) Fraction of successfully localized single-molecules. Time-integrated (25 ms) single-molecule fluorescence signals produce images that resemble PSFs that are blurred to different extents (insets). Faster moving molecules are localized less efficiently due to motion blurring. (c) Expected distributions of apparent diffusion coefficients when confinement and motion blur is taken into account. The similarity of the distributions increase for faster diffusion coefficients. Figure panels a and c are adapted from Ref. (4).

To generate libraries of simulated distributions for arbitrary diffusion coefficients, we performed Monte Carlo simulations of confined Brownian motion inside the volume of a cylinder using a set of 64 diffusion coefficients ranging from 0.05–20 μm^2^/s as input parameters. The size of the confining cylinder was chosen to match the average size of a typical rod-shaped bacterial cell (radius = 0.4 μm, length = 5 μm). The starting position of the trajectory was randomly set within the volume of the cylinder and Brownian motion was simulated using short time intervals of 100 ns. If a molecule was displaced outside of the volume of the cylinder within a time step, it was redirected back towards the inside of the cylinder at a random angle. Choosing a short time step ensured that the entire volume of the cylinder, including the interfacial region near the cell boundary, could be sampled by the diffusing molecule.

To simulate the raw experimental observable, we generated noisy, motion-blurred single-molecule images. For 2D simulations, we summed 50 standard PSFs (approximated as 2D Gaussians with FWHM ~ 325 nm) corresponding to 50 periodically sampled positions of a fluorescent emitter during the camera exposure time (25ms). Similarly, for 3D simulations, we summed 50 DHPSFs. Because the DHPSF has a larger cross section than the standard PSF, fewer photons are necessary for localizing emitters in 2D. To match photon counts measured experimentally, we scaled the photon count of each simulated image to 500 photons per localization for the standard PSF and 1000 photons per localization for the DHPSF. To normalize to the total photon budget, we simulated 3D trajectories with 5 displacements (3D) and 2D trajectories with 11 displacements. To each simulated frame, we added a laser background of ~13 photons/pixel and introduced Poisson noise based on final photon count in each pixel. A dark offset (50 photons/pixel on average) with Gaussian read noise (σ ~1.5 photons) was added as well to produce the final image. The resulting image was then multiplied by the experimentally measured pixel-dependent gain of our sCMOS camera to obtain an image in units of detector counts.

By explicitly simulating spatially blurred emission profiles with realistic signal to-noise ratios, we can account for both static and dynamic localization error. Static localization error is the result of finite numbers of fluorescence signal photons that provide an imprecise measure of the PSF shape and thus result in single-molecule localizations of limited precision (1). Dynamic localization errors manifest for moving emitters that generate motion-blurred images on the detector (**Fig 2b inset**). When analyzed using common fitting algorithms (which are based on data fitting to well-defined PSF shapes), motion-blurred images provide 2D or 3D position estimates with limited accuracy and precision (38). If the motion blur is too severe, then the point-spread-function (PSF) of the molecule may become too distorted to result in a successful fit. Motion blur therefore limits the detection efficiency of fast diffusing molecules (**Fig 2b**).

We simulated *N* = 5000 single-molecule trajectories for each of the 64 input diffusion coefficients to obtain 5000 apparent diffusion coefficient estimates and 5 × 5000 = 25,000 molecular displacements (3D data) or 11 ×5000 = 55,000 molecular displacements (2D data). The corresponding probability density functions *PDF(D*)* and the empirical cumulative distribution functions *CDF(D*)*, or alternatively *PDF(r)* and *CDF(r)*, were then smoothed by B-spline interpolation of order 25 and normalized individually (**Fig. 2c** and **Fig. S1, S2 in the Supporting Material**). The interpolated distributions were then interpolated again along the *D*-axis (*D* is the unconfined (input) diffusion coefficient) using the ‘natural’ interpolation method in the ‘scatteredInterpolant’ MATLAB function. This two-step interpolation provides a continuous function that provides the experimentally expected distribution for any species whose Brownian motion is governed by a diffusion coefficient value in the range of 0.05 and 20 μm^2^/s. The simulated distributions account for the effects of molecular confinement due to the cell boundaries, signal integration over the camera exposure time, and the experimentally calibrated signal-to-noise levels.

### Data Fitting

To estimate the number of diffusive states, their diffusion coefficients, and their population fractions, we fit the experimentally measured cumulative distribution functions using linear combinations of simulated *CDF(r)* or *CDF(D*)*. Using the CDF for fitting instead of a PDF histogram eliminates bin-size ambiguities that can bias the fitting results. To determine the number of diffusive states, we performed a constrained linear least-squares fit (using the ‘lsqlin’ function in MATLAB) and a periodically sampled array of simulated CDFs. We combined diffusive states that had diffusion coefficient values within 20% of each other into a single diffusive state by a weighted average based on their population fractions. The resulting vector of fitting parameters, consisting of diffusion coefficients of individual diffusive states and their respective population fractions, was used as a starting point to create arrays of trial fitting parameter vectors with different numbers of diffusive states, ranging from a single diffusive state to a user-defined maximum number of states (five in all cases considered here). We generated the trial parameter vectors as follows: We either combined adjacent diffusive states through weighted averaging or we split diffusive states into two states with equal population fractions and diffusion coefficient 20% above and below the original value. We considered all state combination and splitting possibilities. We used each trial vector as a starting point for non-linear least-squares fitting of 5 separate subsets of the data (using the ‘fmincon’ function in MATLAB). In each case, the quality of the fit (quantified as the residual sum of squares) was found by comparing the quality of the fit with respect to the remaining subsets (data cross-validation). The average residual sum of squares was used to quantify the quality of the fit corresponding to a given trial vector. This method yielded multiple trial vectors given the number of diffusive states.

For each number of diffusive states, only the trial vector with the best quality of fit was retained. The optimal number of states was then determined by identifying the last trial vector for which adding an additional state resulted in at least a 5% improvement in the quality of the fit. Finally, this trial vector was then used as the starting point to fit the full data set using non-linear least squares fitting. To estimate error in each of the fitted parameters, we resampled the dataset 100 times by bootstrapping and then fit them individually, initializing the fit with the same starting parameter vector. To constrain the optimization, the population fractions of diffusive states below 0.5 μm^2^/s were not refined through non-linear least-squares fitting, but instead assigned to stationary molecules. This choice was made because even completely stationary molecules exhibit non-zero apparent diffusion coefficients in single-molecule tracking experiments due to finite single-molecule localization precision (static localization error). For simplicity, all data and fits are displayed as PDFs instead of CDFs throughout this manuscript.

### Simulation of MINFLUX Trajectories

To simulate experimental tracking data obtained by MINFLUX microscopy, we first computed three-dimensional isotropic Brownian motion trajectories, sampled at high time resolution and confined within a spherocylinder of length *l* = 5 μm and radius *r* = 0.4 μm (same as for camera-based tracking). The short time-step for each displacement was 1 μs and the total trajectory length was 20 ms. We assumed exponentially distributed fluorescence blinking on- and off-times with *t*_*on*_ = 2 ms and *t*_*off*_ = 0.6 ms, in agreement with experimental measurements of the flurorescent protein mEos2 (39). As before, we simulated 5000 trajectories for 64 diffusion coefficients in the range of *D* ∈ [0.05,15] μm^2^/s to create libraries of distributions used for fitting of simulated experimental data. We then projected the 3D motion trajectories onto the *xy*-plane and tracked the blinking emitters using a doughnut intensity profile scanned over the emitter using a 4-step multiplex cycle, as described previously (39). The doughnut size parameter was set to *fwhm* = 800 μm and the field-of-view scanning parameter was set to *L* = 400 μm. Choosing larger values for *fwhm* and *L* minimizes the probability of fast moving emitters (D > 5 μm^2^/s) escaping from the MINFLUX observation region during tracking. The multiplex cycle time was *Δt* = 200 μs. To account for motion blurring during a multiplex cycle, we considered the excitation and emission probabilities from each of the computed emitter positions (sampled at 1 μs time steps). The detected photon counts were assumed to follow Poisson statistics. Emitter localization was performed with the previously described modified least mean squared (mLMS) estimator (39), with *k*=2, *β*_*0*_ = 0.96 and *β*_*1*_ = 5.75. The resulting trajectories each had 100 localizations, which were sampled every 200 μs.

### Modeling State Transition Simulation

To address the effect of a dynamic equilibrium between two diffusive states, we simulated trajectories for which one or more state transitions take place during a single-molecule trajectory. 3D state-switching trajectories were simulated with track lengths of 5 displacements. 2D MINFLUX state-switching trajectories were simulated with track lengths of 99 displacements. We considered a two-state system in which molecules spend equal amounts of time in each state, resulting in a populations fractions of 50% for each state. The average time, *T*, that a molecule takes to switch from one state to the other and back again is

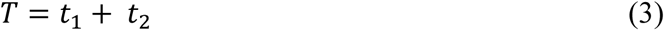

where *t*_*1*_ and *t*_*2*_ are the average time spent in states 1 and 2, respectively. The state-switching kinetics were modeled as follows: Each individual molecule trajectory randomly started in one of the two states. The time *t* spent in a given state before transitioning to the other was modeled as the exponential decay

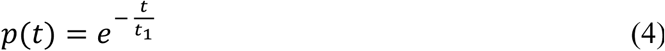

Thus, the time spent in a given state is given by

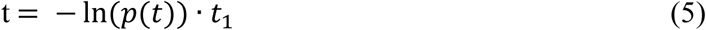

where the value of *p(t)* was a value between 0 and 1 randomly chosen from a uniform distribution. This process was repeated, allowing the molecule to switch back and forth between the two states, until the total amount of time reached the total length of the trajectory. State-switching trajectories were then simulated for camera-based or MINFLUX-based tracking as described above.

### Bacterial Strains and Plasmids

Plasmids for the inducible exogenous expression of fluorescent and fluorescently-tagged proteins were derived from IPTG-inducible pAH12 and arabinose-inducible pBAD vectors. The coding sequences of eYFP were PCR amplified using Q5 DNA polymerase (New England Biolabs, Ipswich, Maine) from pXYFPN-2 (40). The PCR product was isolated using a gel purification kit (Invitrogen, Carlsbad, California) and used as a megaprimer for amplification and introduction into a pAH12-derivative containing a kanamycin resistance cassette, LacI, and a lac promoter to generate pAH12-eYFP. The pAH12 backbone was a gift from Carrie Wilmot.

For the pBAD-mEos3.2, the protein coding sequence was amplified from a mEos3.2-N1 plasmid, gifted to us by Michael Davidson (Addgene plasmid # 54525). The PCR products were gel purified, and both the PCR products and the pBAD-backbone were digested with EcoRI and XhoI restriction enzymes (New England Biolabs). Digested vector and inserts were ligated using T4 DNA ligase and transformed into *E. coli* TOP10 cells. Colonies were PCR screened for presence of correct insert using GoTaq DNA Polymerase (Fisher Scientific, Hampton, New Hampshire), and plasmid was isolated from positive clones (Omega Biotek, Norcross, Georgia)

All plasmids were sequenced by GeneWiz (South Plainfield, New Jersey) prior to electroporation into *Y. enterocolitica* for analysis. Transformed cells were plated on LB agar [10 g/L peptone, 5 g/L yeast extract, 10 g/L NaCl, 1.5% agar] (Fisher Scientific, Hampton, New Hampshire) containing kanamycin [50 μg/mL] or ampicillin [200 μg/mL]. For electroporation of *Y. enterocolitica* pIML421asd cells, recovery media and plates also contained diaminopimelic acid (dap). A list of all strains and plasmids can be found in **Table S1 in the Supporting Material**.

### Cell Culture

*Y. enterocolitica* cultures were inoculated from a freezer stock in BHI media (Sigma Aldrich, St. Louis, Missouri) with nalidixic acid (Sigma Aldrich) [35 μg/mL] and 2,6- diaminopimelic acid (Chem Impex International, Wood Dale, Illinois) [80 μg/mL] one day prior to an experiment and grown at 28°C with shaking. After 24 hours, 300 μL of overnight culture was diluted in 5 mL fresh BHI, nalidixic acid, and diaminopimelic acid (dap) and grown at 28°C for another 60-90 minutes. In addition, inoculation media also contained kanamycin or ampicillin for pAH12- or pBAD-based plasmids, respectively. Cultures of cells containing pAH12- or pBAD-based plasmids were induced with IPTG (Sigma Aldrich) [0.2 mM, final] or arabinose (Chem Impex) [0.2%], respectively, for the final 2 hours of incubation. Cells were pelleted by centrifugation at 5000 g for 3 minutes and washed 3 times with M2G (4.9 mM Na_2_HPO_4_, 3.1 mM KH_2_PO_4_, 7.5 mM NH_4_Cl, 0.5 mM MgSO_4_, 10 μM FeSO4 (EDTA chelate; Sigma), 0.5 mM CaCl_2_) with 0.2% glucose as the sole carbon source). The remaining pellet was then re-suspended in M2G and dap. Cells were plated on 1.5 – 2% agarose pads in M2G containing dap.

## Results and Discussion

### eYFP and mEos3.2 undergo confined Brownian Diffusion in *Y. enterocolitica*

To experimentally validate the numerical analysis framework based on Monte Carlo simulations of confined diffusion, we tracked the 3D motion of individual eYFP and mEos3.2 fluorescent proteins in living *Y. enterocolitica* cells. Previous studies in *E. coli* (28, 41) and *C. crescentus* (42) have established that small cytosolic proteins undergo Brownian motion. Non-specific interactions due to macromolecular crowding reduce the diffusion coefficient for small cytosolic proteins, but do not by themselves lead to measurable deviations from normal Brownian diffusion (43). In contrast, the motion of large macromolecular complexes (>30 nm in diameter) is best described by anomalous diffusion due to glass-like properties of the bacterial cytoplasm (44).

The experimentally measured distributions of apparent diffusion coefficients are fit well using a single diffusive state with *D* = 11.3 μm^2^/s (for eYFP, **Fig. 3a**) and *D* = 15.0 μm^2^/s (for mEos3.2, **Fig. 3b**). The close agreement between simulations and experiment confirms that the assumption of spatially confined Brownian diffusion is valid for both eYFP and mEos3.2 in *Y. enterocolitica* under our experimental conditions. These diffusion coefficient values are in agreement with previously measured values of GFP in bacteria (28, 45-50). The structure and molecular weights of eYFP (27 kDa) and mEos3.2 (26 kDa) are very similar. The differences in their diffusion coefficients may thus be due to differences in non-specific transient interactions with other cellular components. We also note that there is a small (6% or less) stationary (<0.5 μm^2^/s) population for both fluorescence proteins. We find small numbers of stationary trajectories in all of our single-molecule tracking datasets, which indicates that even freely diffusing cytosolic proteins may become immobilized. However we did not find that that these stationary molecules exhibit any subcellular preference.

**Figure 3.**
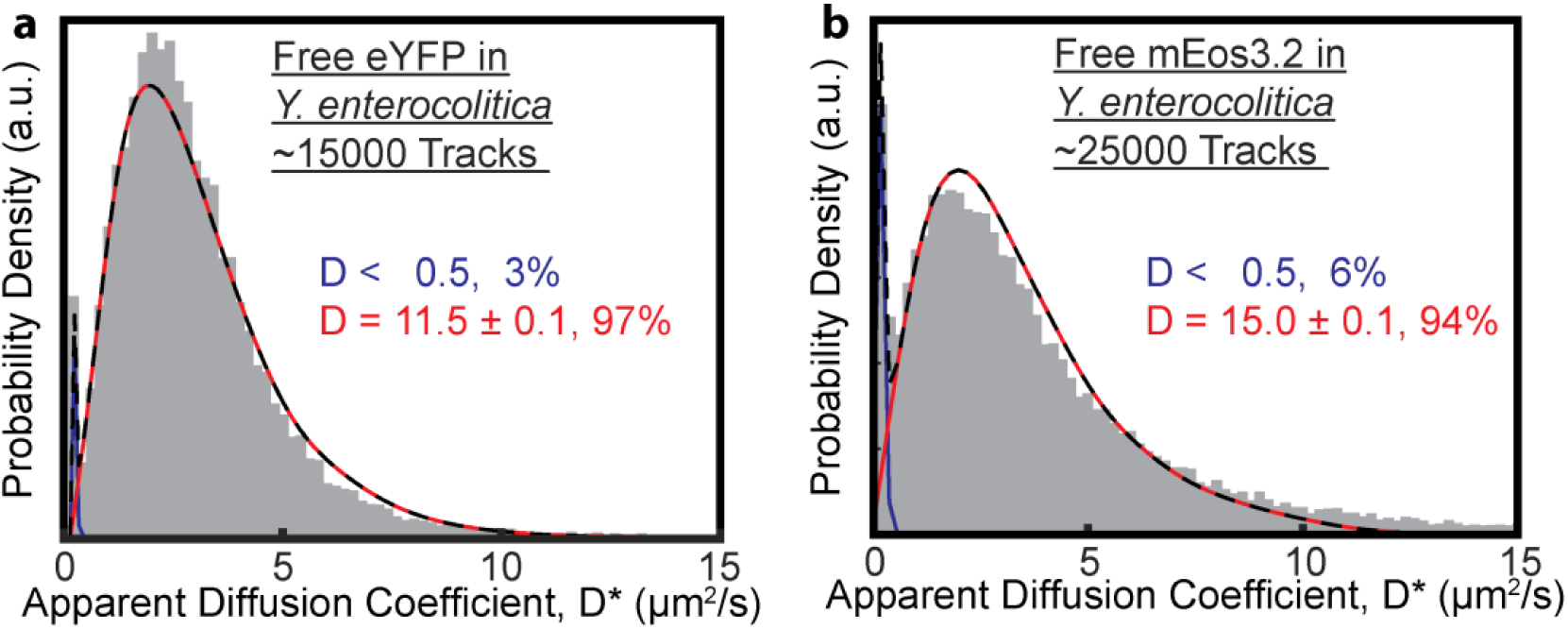
The 3D diffusion of cytosolic fluorescent proteins eYFP and mEos3.2 in *Y. enterocolitica* can be explained using a single diffusive state. (a) eYFP diffuses at 11.5 μm^2^/s (red). (b) mEos3.2 diffuses at 15.0 μm^2^/s (red). A small fraction (<6%) of stationary trajectories is present in both datasets (blue). The total fit is shown as a dashed black line.

### 2D vs 3D Single-Molecule Tracking to Estimate Diffusion Coefficients

Most single-molecule tracking results reported to-date utilize the standard PSF for 2D single-molecule tracking. Acquiring 3D trajectories requires engineered PSFs, such as astigmatic, double-helix, or tetra-pod PSFs (10, 51-55). A common feature of engineered PSFs is their increased footprint on the detector compared to the standard PSF. Due to their increased size, engineered PSFs require higher photon counts to achieve lateral localization precisions equivalent to those obtained with the standard PSF. Given the finite photon-budgets of fluorescent labels, 2D tracking can thus yield longer single-molecule trajectories that contain roughly twice the number of displacements than 3D trajectories acquired with engineered PSFs.

To determine whether diffusion coefficients are more accurately estimated by 2D or by 3D tracking, we repeated the 3D DHPSF simulations using the standard PSF. We generated simulated distributions of apparent 2D diffusion coefficients in the same way as for the 3D data (Materials and Methods). However, the simulated 2D trajectories had twice as many displacements as the 3D trajectories to provide an equivalent total photon count over the course of a trajectory. We found that the resulting 2D apparent diffusion coefficient distributions are broader and their peaks are systematically right-shifted compared to their 3D equivalents (**Fig. 4a**). The increased left-shift of the 3D distribution is due to the additional confinement of the molecule’s motion in the *z*-dimension that is not measured in 2D tracking.

**Figure 4.**
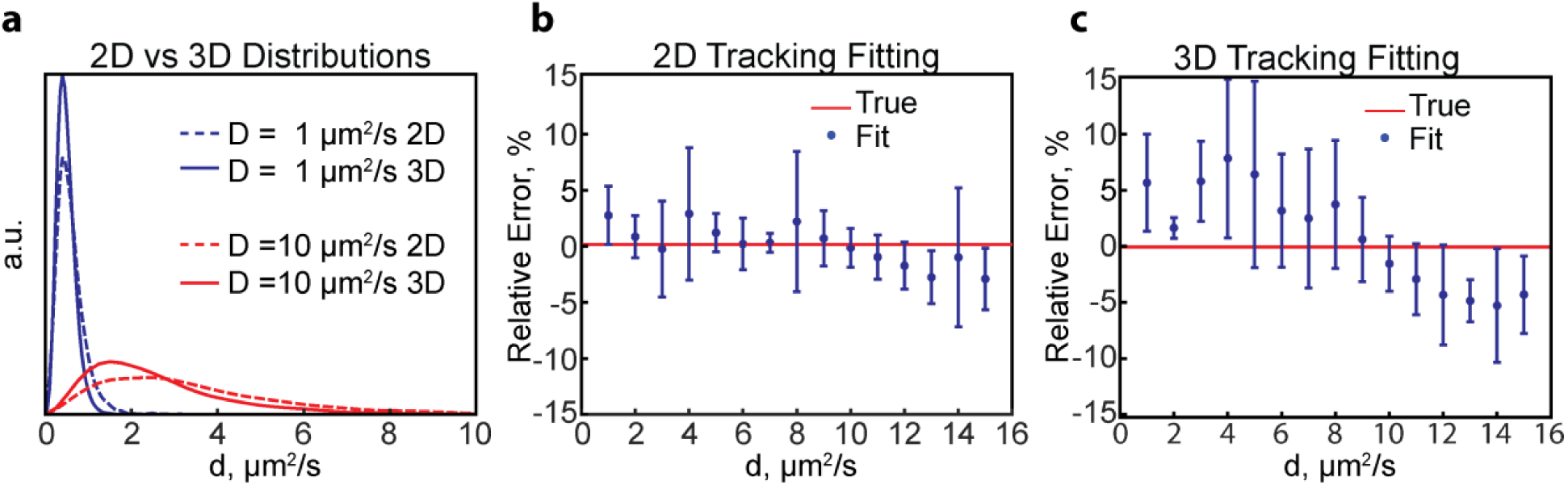
Comparison of 2D and 3D tracking. (a) Comparison of 2D and 3D apparent diffusion coefficient distributions corresponding to 1 μm^2^/s and 10 μm^2^/s. The distributions for 3D tracking are left-shifted to a larger extent due to the additional confinement in the 3^rd^ dimension. (b,c) Relative errors in determining the diffusion coefficient of a single diffusive state using 2D (b) and 3D (c) single-molecule tracking. Shown are the averages and standard deviations of four independent simulations containing *N* = 5000 trajectories each resampled 10 times by bootstrapping.

We then performed numerical fitting of simulated 2D tracking data to estimate the diffusion coefficient. We found that there is a slight increase in accuracy when fitting 2D data compared to 3D data for a single diffusive state, particularly for fast diffusion. (**Fig. 4b,c**). The improved accuracy of 2D tracking may be due to the decreased similarity of the 2D distributions for fast diffusion coefficients (**Fig. S1**), which enables more accurate parameter estimation.

### Single-molecule tracking can be used to resolve different diffusive states

The free fluorescent proteins examined in the previous section each exhibited a single predominant diffusive state, which means that these two proteins do not exhibit stable interactions with other cellular components. This property is important for their use as non-perturbative labels that do not alter the diffusive behaviors of the target proteins beyond an overall reduction in their native diffusion rate. An overall reduction in diffusion rate is expected due to the increased molecular weight and hydrodynamic radius of the fusion protein. If the target protein stably interacts with cognate binding partners to form homo- or heterooligomeric complexes of different sizes, then single-molecule tracking of non-perturbatively labeled target proteins may be used to resolve the corresponding diffusive states. Examples of different diffusive states reported in the recent literature include the cytosolic pre-assembly of the bacterial type 3 secretion system proteins SctQ and SctL (4), ternary complex formation of the elongation factor Tu (EF-Tu) which can bind to aminoacyl-tRNA, GTP, and translating ribosomes(17), the nucleotide excision repair initiation molecule UvrB (14), and short-lived ribosome binding of EF-P(13).

To test the resolving capability of single-molecule tracking, we simulated mixed distributions of 3D displacements or apparent diffusion coefficients that contain two different diffusive states. We then fit these distributions to obtain the unconfined diffusion coefficients and relative population fractions of each diffusive state. By systematically varying the diffusion coefficients, we assessed the error in the optimized fitting parameters for various combinations. We examined both equal (50:50) and unequal population fractions (80:20). In all cases, the distributions were based on 5000 trajectories with five displacements each. We found that the errors in the optimized fitting parameters increased when the diffusion coefficients were similar, as evidenced by the wedge-shaped diagonal (**Fig. 5a**). Slight differences in diffusion rate are thus more readily resolved for slowly diffusing molecules than for faster moving ones. We reason that the ability to resolve fast diffusive states is further compromised by the confinement effect, which causes the distributions of apparent diffusion coefficients to become more similar in the high diffusion coefficient limit (**Fig. 2c**).

**Figure 5.**
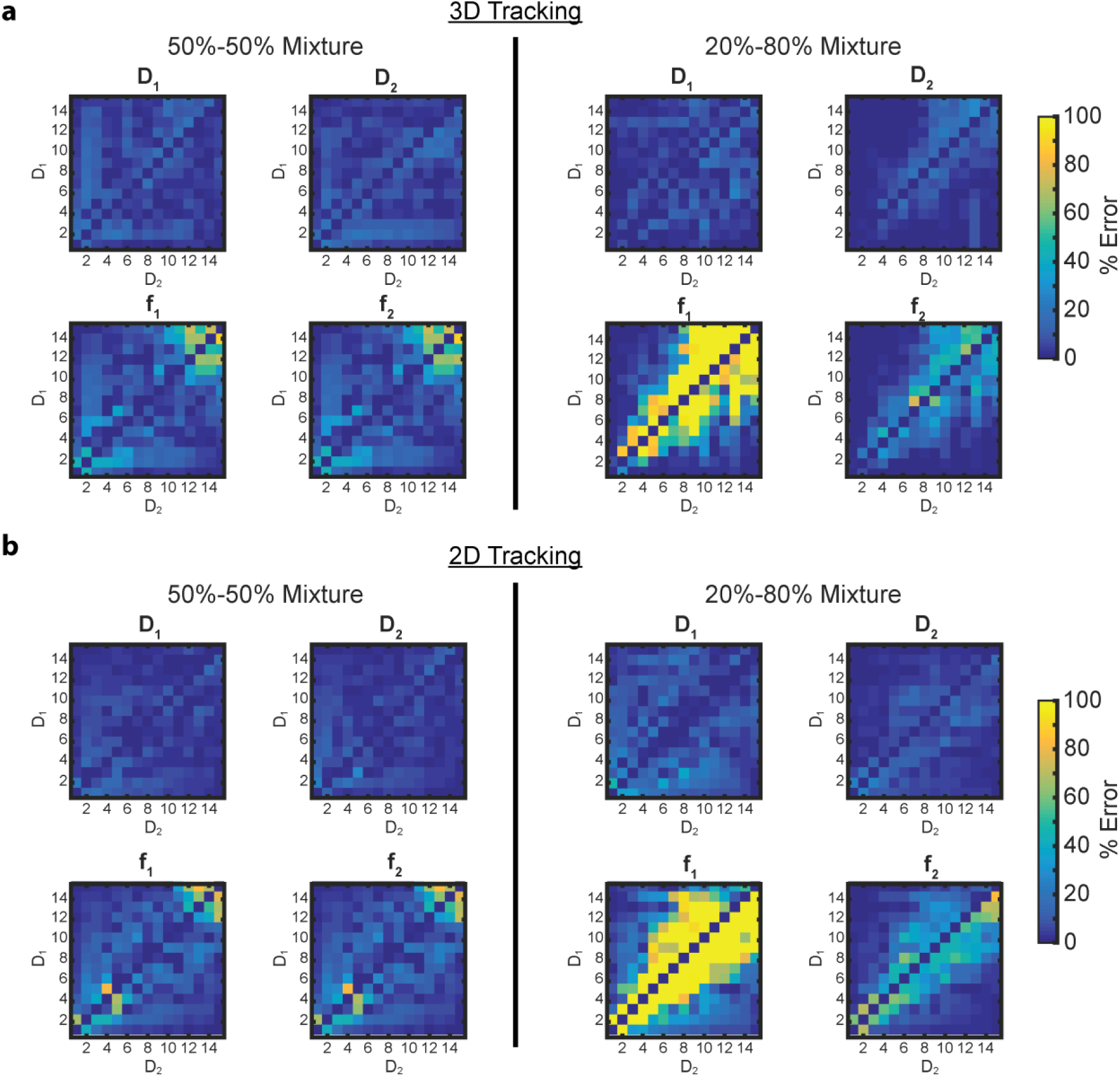
Multiple diffusive states can be resolved by numerical fitting of single-molecule tracking data using 2D and 3D tracking. (a,b) Relative errors for determining the diffusion coefficients and population fractions of binary mixtures of diffusive states using 3D (a) and 2D (b) tracking. The relative population fractions in the two state mixtures were either 50%-50% (left) or 20%-80% (right). The relative error for each fitting parameter (diffusion coefficients D1 and D2, and their corresponding population fractions f_1_ and f_2_) is represented as a matrix for different diffusion coefficient combinations. Each pixel represents the mean (relative) error of the parameter’s fit value after analyzing ten datasets (resampled by bootstrapping) each containing 5000 tracks.

Current detector technologies, in particular large field-of-view sCMOS detectors, have made it possible to readily acquire single-molecule trajectories in thousands of cells in a single imaging session. Thus, 5,000 trajectories can be obtained even for proteins expressed at low levels. For highly expressed proteins up to 100,000 trajectories can be obtained. We therefore repeated our analysis using distributions based on 100,000 trajectories. As expected, the errors in the parameter estimates decreased (~7% on average) when fitting the now more thoroughly-sampled distributions (**Fig. S3 in the Supporting Material**). Therefore, the resolving capability improves when additional measurements are available to sample the shape of experimental distributions. However, larger errors persist along the diagonal of the error matrices, highlighting the difficulty in resolving states with similar diffusion coefficients. When the population fractions are split 80:20, larger errors manifest due to the smaller number of proteins in the diffusive state with a 20% population fraction. In those cases, the relative error in the smaller fraction can approach 100%, i.e. the smaller fraction is completely eliminated when the fitting routine converges on a one-state solution (**Materials and Methods**).

To test whether the above results may be extrapolated to more complex state distributions, we simulated a few selected examples of mixed distributions containing three and four diffusive states, maintaining *N* = 5000 total trajectories in each case. We found that three states can be simultaneously resolved as long as their diffusion coefficients are sufficiently different and their population fractions are similar (**Fig. S4a in the Supporting Material**). Again, the errors in the fitting parameters increase for faster (i.e. more similar) diffusion coefficients (**Fig. S4b**). In the case of a 4-state population, the distribution is best fit with a 3-state results, even when the values of the diffusion coefficients are well separated (**Fig. S4c**). Specifically, the two fastest states are combined into a single state with a correspondingly larger population fraction. The 3- and 4- state simulations thus recapitulate the trends observed for binary diffusive state mixtures.

To test whether 2D tracking is also more discriminating when multiple diffusive states are present, we constructed simulated 2-state distributions of apparent diffusion coefficients based on 2D data. Again, we observed only a slight increase in the accuracy of the fitting (~3%) for the 2D fitting compared to 3D for a two state fitting **(Fig 5b**). We therefore conclude that 2D and 3D single-molecule tracking are roughly equivalent in their ability to resolve different diffusive states. We note however that 3D single-molecule localization microscopy has the additional advantage of providing more detailed spatial information on the subcellular locations of diffusing molecules, which may provide important additional information in select cases. We also note that the above analysis only pertains to diffusion of cytosolic proteins. The diffusion of membrane proteins is subject to different confinement effects that may make it more appropriate to track in 3D (5).

### Transitions between diffusive states

Thus far, we have only considered diffusive states that do not interconvert on the time-scale of a single-molecule trajectory (~100-300 ms on average). Under physiological conditions, however, molecules may frequently bind to or dissociate from cognate interaction partners and thereby transition between different diffusive states. The time-resolution for making single-step displacement measurements (~25 ms) is shorter than the time resolution for determining apparent diffusion coefficients (~5 · 25 ms = ~125 ms). We therefore hypothesized that, in the presence of diffusive state switching, more accurate parameter estimates may be obtained by fitting single-step displacement distributions. To test this hypothesis, we simulated distributions for two states, D_1_ = 1 μm^2^/s and D2 = 10 μm^2^/s, that can interconvert on timescales comparable to a single-molecule trajectory. We then gradually decreased the average diffusive state switching time *T* = (*k*_*1*_)^−1^ + (*k*_*1*_)^-1^ = *t*_*1*_ + *t*^*2*^ and imposed *k*_*1*_ = *k*_*2*_ to keep the population fractions equal (**Materials and Methods**). To fit the single-step displacement distributions, we generated a library of simulated single-step displacement distributions as described before for apparent diffusion coefficients (**Fig. S2**). Both the apparent diffusion coefficient distributions and single-step displacement distributions were then fit with their respective library. To quantify the overall accuracy of the fit, we averaged the relative errors of all fitting parameters (in this case the diffusion coefficients *D*_*1*_ and *D*_*2*_ and the population fractions *f*_*1*_ and *f*_*2*_ = 1 – *f*_*1*_. We found that, in the limit of infinitely long switching times (no state transitions), both approaches produce parameter estimates with similar accuracy (**Fig. 6a,b** and **Fig. S5 in the Supporting Material**). As the average switching time is decreased, the mean relative errors start to increase for both methods. Importantly, fitting distributions of apparent diffusion coefficients produced parameter estimates that deviated sooner from the ground truth (as a function of decreasing average switching time) than those obtained by fitting single-step displacement distributions. In the limit of short switching times, fitting of both the apparent diffusion coefficient and single-step displacement distributions produced large errors, because a single molecule can sample both diffusive states repeatedly during the timescale of the measurement. When using 25 ms exposure times, accurate parameter estimates can be made for this two-state system, if *T* > 75 ms and *T* > 500 ms for displacement and apparent diffusion coefficient fitting, respectively. For accurate extraction of the parameters, the time resolution of the measurement should be about three times shorter than the average switching time *T*.

**Figure 6.**
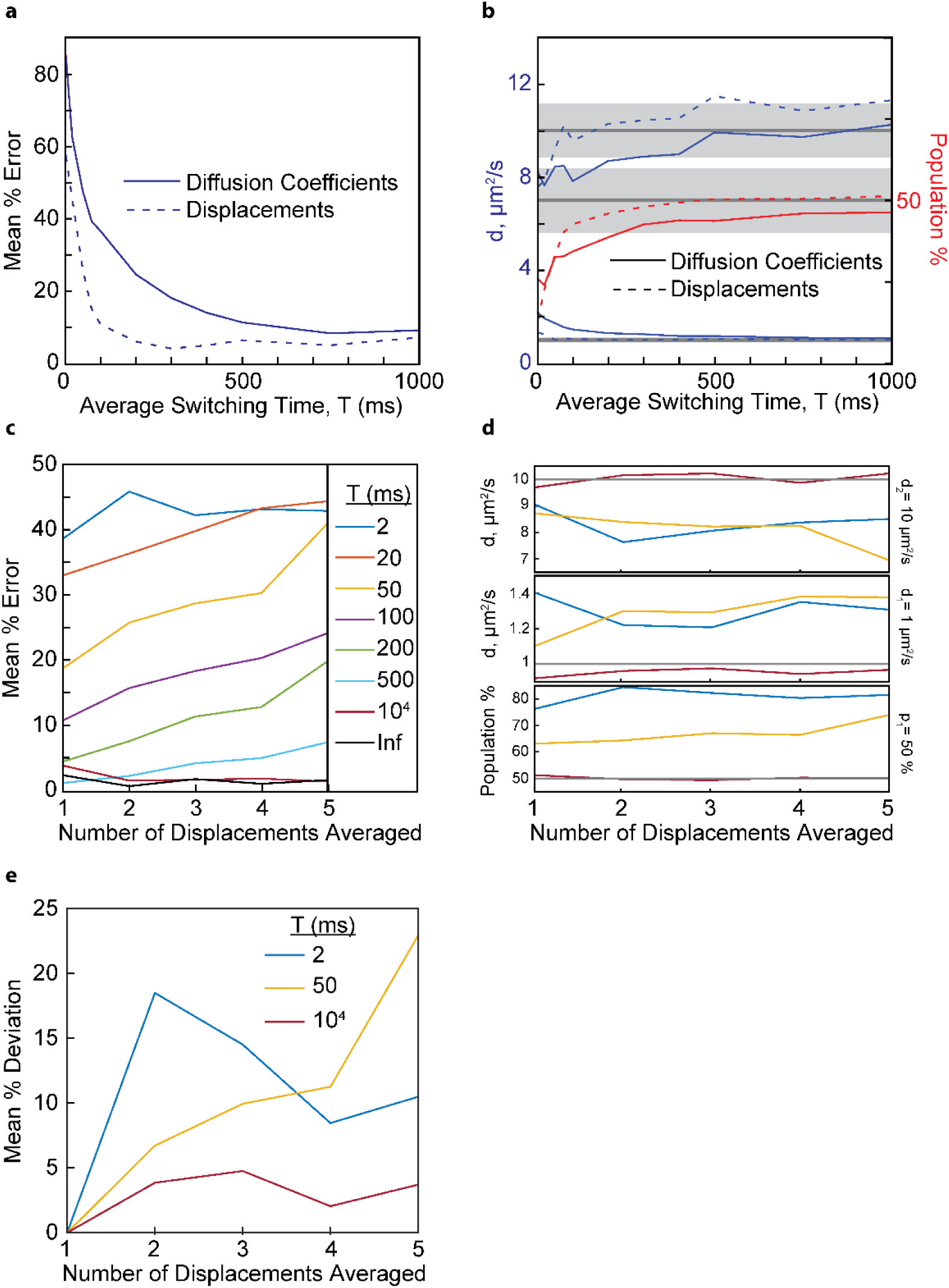
Resolving diffusive states in the presence of dynamic state transitions. (a) The mean relative errors of the fitting parameters for a 2-state mixture (D_1_ = 1 μm^2^/s, D_2_ = 10 μm^2^/s, 50:50 population fraction) as a function of different switching times between two diffusive states. The mean % error obtained by fitting the single-step displacement distributions diverges for *T* < 75 ms, whereas the mean % error obtained by apparent diffusion coefficient fitting diverges for *T* < 500 ms. (b) Individual parameter estimates as a function of state switching time for the same simulations as in (a). Population fraction *f*_*2*_ = 1 – *f*_*1*_ is not shown for clarity. (c) The mean relative errors of the fitting parameters as a function of the number of averaged displacements. The shaded areas represent 10% error limits for each parameter. (d) Parameter estimates as a function of averaged displacements for the same simulations as in panel c. Color scheme is the same as the legend in panel c. Grey lines represent the ground truth. The fitted individual parameter value produces horizontal curves for both the very short (2 ms) and very long (10^4^ ms) switching times. For intermediate switching times (50 ms), the fitted values trend away from the true value as the number of averaged displacements increases. (e) Mean deviation relative to the single displacement parameter estimates (*N*_*i*_ = 2) for different switching times.

The above observations suggest that it should be possible to estimate the timescale of diffusive state switching by time-averaged diffusion (TAD) analysis, i.e. by varying the number of averaged displacements. We therefore evaluated the apparent diffusion coefficients for overlapping sub-trajectories having different numbers of displacements/localizations. Specifically, within each single-molecule trajectory, we define overlapping sub-trajectories with *N*_*i*_ localizations and *N*_*i*_ *-1* displacements. The number of sub-trajectories for a given *N*_*i*_ is *S=N-N*_*i*_*+1*, where *N* is the number of localizations in the full-length trajectory. Defining the first localization in the sub-trajectories as *P*, we modified Eqn 1 to

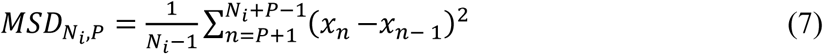

to obtain mean squared displacement values for different sub-trajectory lengths and starting points, namely *N*_*i*_ = 2, 3, …, 6 and *P=1…S.*

Based on these sets of observables, we generated five new apparent diffusion coefficient libraries corresponding to the five different values of *N*_*i*_ (on average our experimental 3D trajectories are 5 displacements long). The state-switching trajectories were then re-analyzed using Eqn 7 and fit with the corresponding library. Again, we used the mean relative error over all fitting parameters to quantify the overall accuracy of the fit for each value of *N*_*i*_ (**Fig. 6c**). Consistent with the results above, the accuracy of the fitting parameters is poor for short switching times and good for long switching times. Importantly, the mean relative errors are constant for all *N*_*i*_ in both of these limiting cases. Thus, if the state switching time is substantially shorter or longer than the time resolution of the measurement, then the mean error does not change. In contrast, the mean errors increase for increasing *N*_*i*_, if switching times are comparable to the timescale of a single-molecules trajectory (0.05-0.5s). The same trends are also observed when plotting the individual parameter fitting results (**Fig. 6d**). Based on these results, we conclude that the timescale of diffusive state switching can be estimated by determining the rate of change of individual fitting parameters as a function of the number of averaged displacements. For example, based on the results in **Fig. 6cd**, observing a consistent increase or decrease of individual fitting parameters as a function of *N*_*i*_ would indicate a diffusive state switching time between 20 and 500 ms. We note that the ground truth is unknowable in experimental work. We therefore computed an error relative to the parameter values obtained when fitting single displacement distributions (i.e. *N*_*i*_ = 2). Single displacement distributions offer the best time resolution and thus should be least affected by diffusive state averaging. The parameter deviations relative to the parameter estimates at *N*_*i*_ = 2 displayed similar trends as those referenced to the ground truth (**Fig. 6e**).

It is clear that the dynamic range of TAD analysis improves if trajectories contain a large number of displacements. However, in camera-based tracking of fluorescent fusion proteins, only *N* = 5 or *N* =12 displacements can be observed on average for 3D and 2D tracking, respectively. Longer trajectories can be acquired using chemical dyes (24, 56, 57) or multiple fluorophores as labels (58), but potential of non-specific labeling or the size of multivalent fluorescent tags have to be weighed against this benefit. An important advantage of camera-based tracking is that the temporal dynamic range is tunable to access slow switching timescales (>500 ms) by adjusting the exposure time and/or by acquiring single-molecule trajectories in time-lapse mode (17, 27, 59). On the other hand, exposure times shorter than a few milliseconds come at the expense of data acquisition throughput, because the full chip of current sCMOS cameras cannot be read out faster than 100 Hz (17). Thus, faster timescales are difficult to assess by camera-based tracking.

A solution to access faster time scales is MINFLUX microscopy (39). The time resolution of MINFLUX-based single-molecule tracking is two orders of magnitude better than camera-based tracking (0.2 ms vs 25 ms) and the number of localizations *N* is larger by one order of magnitude (*N*~100 vs. *N*~10). MINFLUX microscopy may thus be able to provide access to state switching dynamics on 0.2 ms to 20 ms timescales, whereas camera-based tracking can cover state switching dynamics on millisecond to minute timescales. To test the capability of MINFLUX microscopy to quantify fast state switching times, we applied TAD analysis to simulated MINFLUX data. MINFLUX trajectories were generated in the same way as the camera-based trajectories, i.e. through Monte Carlo simulations of confined Brownian diffusion, but the MINFLUX localization algorithm was used instead of PSF fitting (**Materials and Methods**). We then used libraries of *Ni*-fold averaged MINFLUX displacement distributions to fit state-switching trajectories for different switching times *T* (*D*_*1*_ = 1 μm^2^/s, *D*_*2*_ = 10 μm^2^/s, *k1* = *k2*). We found that the mean % error vs. *N*_*i*_ curves (**Fig. 7a**) displayed two key characteristics that correlate linearly with switching time *T* or with switching rate 1/*T*. First, for each switching time *T*, there exists a threshold value *N*_*i,T*_, after which the mean % error increases linearly as a function of *N*_*i*_. *N*_*i,T*_ and *T* are linearly correlated (**Fig. 7ab**). Second, the slope of the initial linear increase and the switching rate 1/*T* are linearly correlated as well (**Fig 7ac**). Based on these linear relationships, we conclude that the timescale of state transitions can be determined from the position of *N*_*i,T*_ and from the slope of the following linear increase.

**Figure 7.**
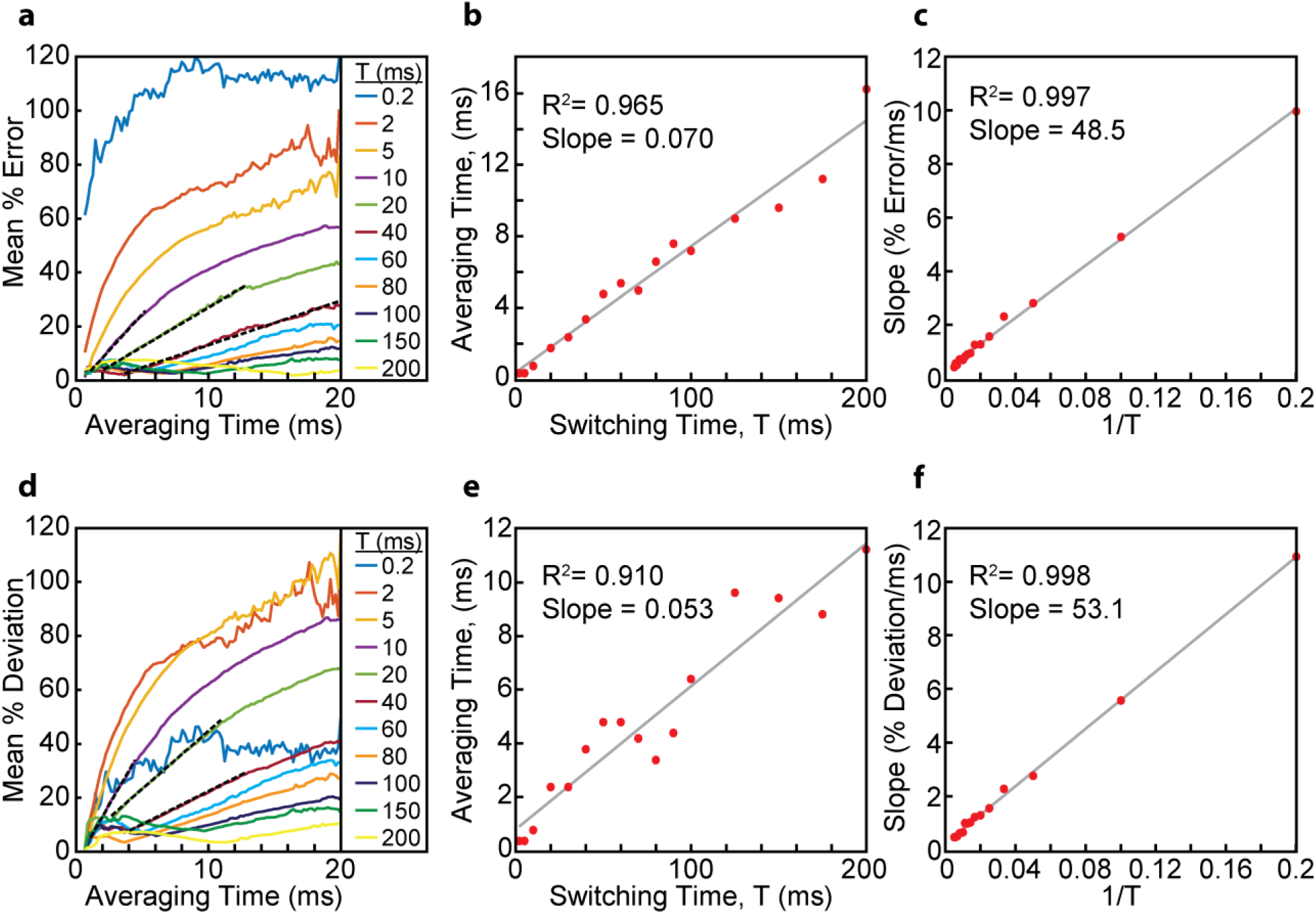
Resolving diffusive states in the presence of dynamic state transitions for MINFLUX data. (a) Mean % error in the parameter estimates compared to the ground truth for various switching times (*D*_*1*_ = 1 μm^2^/s, *D*_*2*_ = 10 μm^2^/s, *k*_*1*_ = *k*_*2*_). Initial slope determinations (dashed black lines) are shown for the *T* = 10, 20, and 40 ms datasets. The averaging time is the value of *N*_*i*_ multiplied by the multiplex cycle time *Δt* = 200 μs. (b) Averaging time at which the mean % error begins to linearly increase. (c) Slope of the initial linear increase of the mean % error. Switching times of 0.2 and 2 ms are not included here, because the linear section of their curves in panel are not sufficiently resolved. (d) Mean % deviation in the parameter estimates relative to the parameter estimates at *N*_*i*_ = 3. Again, initial slope determinations (dashed black lines) are shown for the *T* = 10, 20, and 40 ms datasets. (e) Averaging time at which the mean % deviation in panel d begins to linearly increase. (f) Slope of the initial linear increase of the mean % deviation in panel d. Again, switching times of 0.2 and 2 ms are not included.

Since the ground truth is not accessible by experiment, we repeated the above analysis by referencing all parameter estimates to the parameters obtained at *N*_*i*_ = 3 (**Fig. 7d**). *N*_*i*_ = 3 corresponds to a time resolution of 600 μs. The curves obtained by plotting the mean % deviation from the *N*_*i*_ = 3 parameter estimates vs. *N*_*i*_ displayed the same characteristic linear increases as a function of *N*_*i*_. The onset of the linear increase *N*_*i,T*_ and the slope of the linear increase still correlated linearly with *T* and 1/*T*, respectively (**Fig. 7def**). These results show that the switching rate between two diffusive states can be reliably determined by TAD analysis of 2D and 3D single-molecule tracking data.

## Conclusions

In this work, we present and test a robust analysis method for estimating diffusive state parameters of fluorescently labeled biomolecules in confined bacterial cell volumes based on single-molecule tracking. We show that it is possible to resolve the unconfined diffusion coefficients and the population fractions of multiple diffusive states based on a few thousand short single-molecule trajectories obtained by camera-based tracking. The numerical analysis framework presented is generally applicable to both 2D and 3D tracking and any confinement geometry. We show that 2D and 3D single-molecule tracking are roughly equivalent in their ability to resolve multiple diffusive states. To address the issue of diffusive state switching during the timescale of measurement, we propose time-averaged diffusion (TAD) analysis. By averaging over different number of subsequent displacements, the timescale of state switching can be determined, if that timescale is comparable to the duration of the recorded trajectories. For example, MINFLUX microscopy can provide access to state switching dynamics occurring on 2-200 ms timescales using data acquisition parameters relevant for fluorescent protein localization in living cells. On the other hand, camera-based tracking can be used to detect state switching dynamics on 20 ms to seconds timescales either by using longer exposure times or by acquiring data in time-lapse mode. TAD analysis of experimental single-molecule trajectories thus provides a general and robust approach to quantify the diffusive states and diffusive state transitions that manifest in living cells.

## Supporting information

## Acknowledgements

We thank Dave Cafiso for critical reading and comments on the manuscript. Funding for this work was provided by start-up funds from the University of Virginia.

