## Supplementary material for "Resolving Cytosolic Diffusive States in Bacteria by Single-Molecule Tracking"

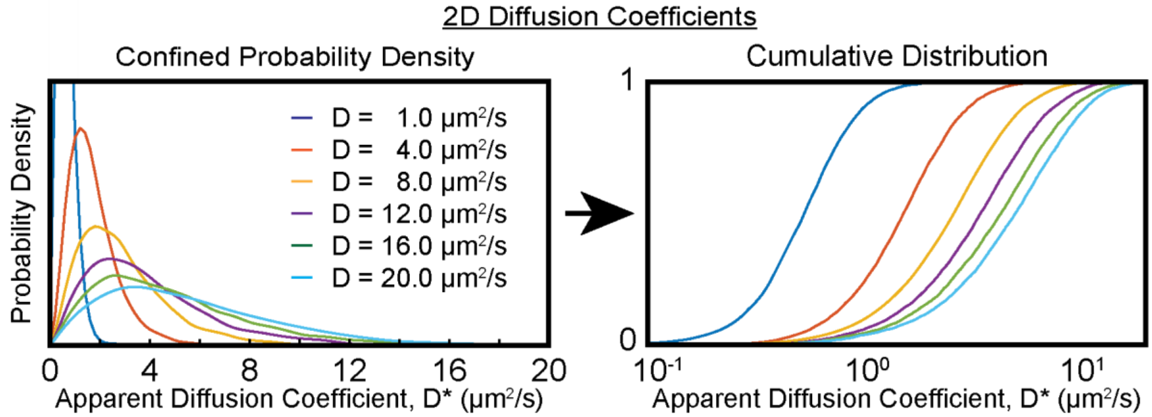


**Fig S1**. Examples of the apparent diffusion coefficient distribution library for 2D tracking equivalent to the 3D distributions shown in Fig. 2b.

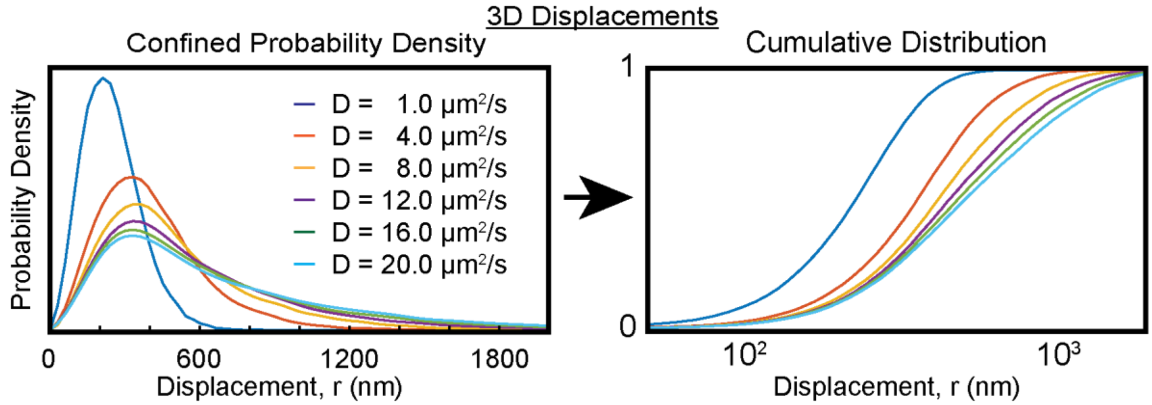


**Fig S2**. Examples of the displacement distribution library for 3D displacements equivalent to the 3D apparent diffusion coefficient distributions shown in Fig. 2b.

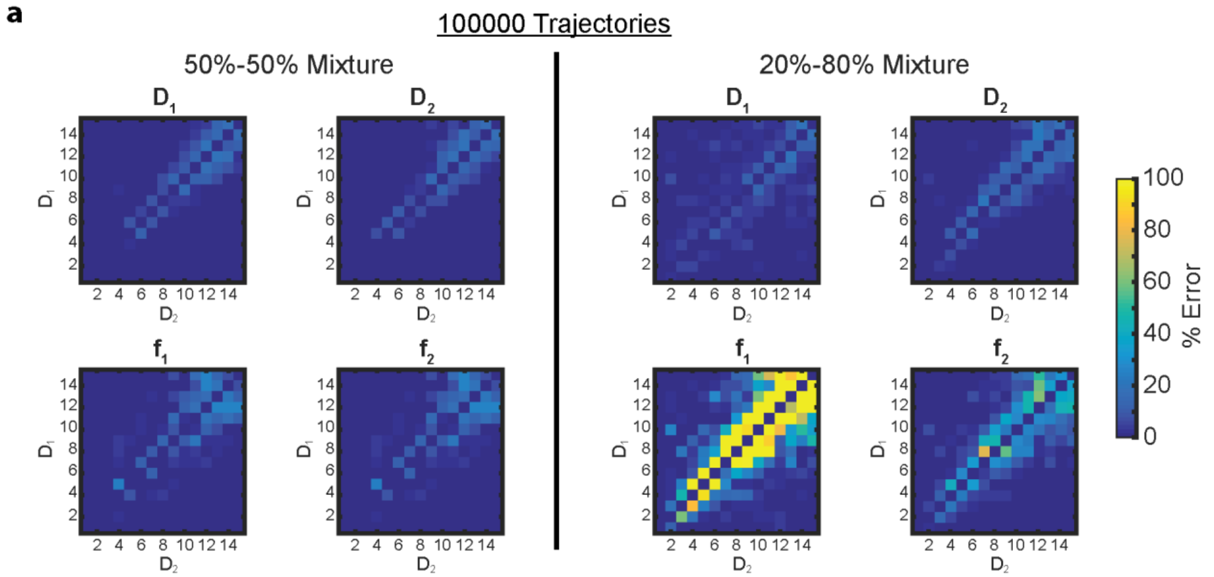


**Fig S3**. 3D apparent diffusion coefficient distribution fitting of 2-state populations with 100000 trajectories for 50%-50% population fraction mixtures (left) or 20%-80% population fraction mixtures (right). The relative errors decrease compared to Fig 5a when the distributions are better sampled with more trajectories.

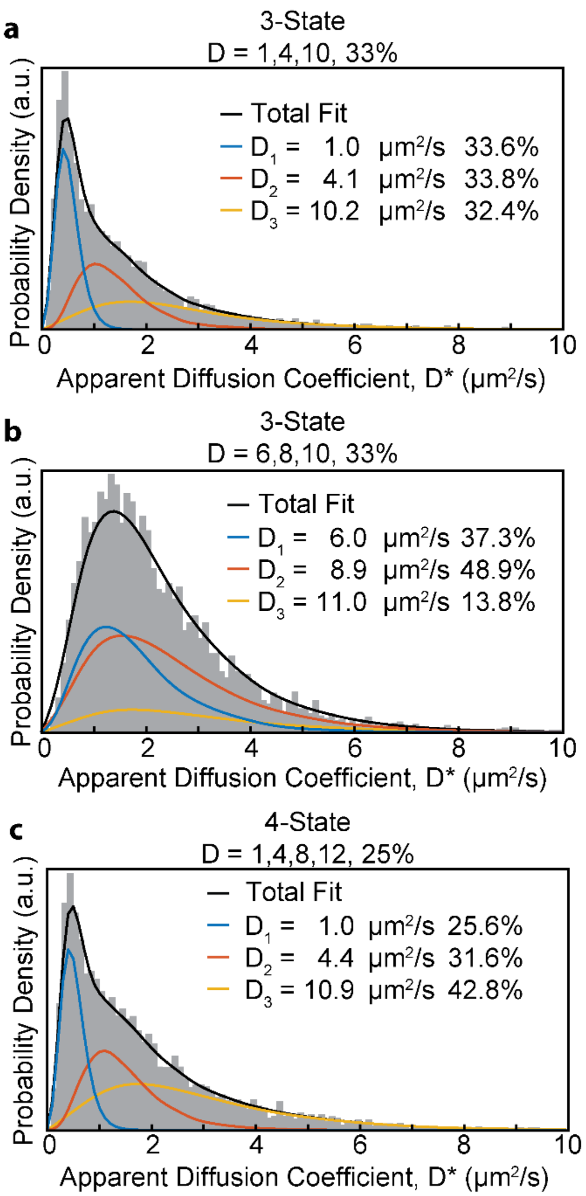


**Fig S4**. Multiple diffusive state fitting examples. (a,b) 3-state distributions were simulated with equal population fractions of 33% each. In (a) the diffusion coefficients are well separated, and both the diffusion coefficients and respective population fractions were accurately fit. However, in (b) the diffusion coefficients were close in value, leading to increased error in the fitting parameters. (c) A 4-state distribution was simulated with equal population fractions of 25% each. The fitting algorithm determined that the best fit was with a 3-state mixture. In this case, the fastest two diffusive states were combined into one state with an intermediate diffusion coefficient.

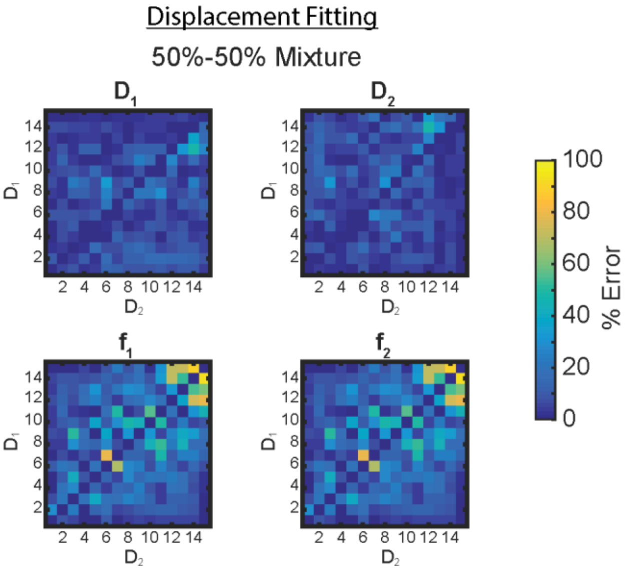


**Fig S5**. 3D displacement distribution fitting of 2-state populations.

**Table S1.** List of strains and plasmids.

| Strain Name | Characteristics | Ref |
|---|---|---|
| AG0003 | pAH12-eYFP | (4) |
| AG0006 | pBAD-mEos3.2 | This work |
